# Waves of Colonization and Gene Flow in a Great Speciator

**DOI:** 10.1101/2024.07.18.603796

**Authors:** Ethan F. Gyllenhaal, Serina S. Brady, Lucas H. DeCicco, Alivereti Naikatini, Paul M. Hime, Joseph D. Manthey, John Kelly, Robert G. Moyle, Michael J. Andersen

## Abstract

Secondary contact between previously allopatric lineages offers a test of reproductive isolating mechanisms that may have accrued in isolation. Such instances of contact can produce stable hybrid zones—where reproductive isolation can further develop via reinforcement or phenotypic displacement—or result in the lineages merging. Ongoing secondary contact is most visible in continental systems, where steady input from parental taxa can occur readily. In oceanic island systems, however, secondary contact between closely related species of birds is relatively rare. When observed on sufficiently small islands, relative to population size, secondary contact likely represents a recent phenomenon. Here, we examine the dynamics of a group of birds whose apparent widespread hybridization influenced Ernst Mayr’s foundational work on allopatric speciation: the whistlers of Fiji (Aves: *Pachycephala*). We demonstrate two clear instances of secondary contact within the Fijian archipelago, one resulting in a hybrid zone on a larger island, and the other resulting in a wholly admixed population on a smaller, adjacent island. We leveraged low genome-wide divergence in the hybrid zone to pinpoint a single genomic region associated with observed phenotypic differences. We use genomic data to present a new hypothesis that emphasizes rapid plumage evolution and post-divergence gene flow.

---

Terrestrial island systems have long served as natural laboratories for studying the process of allopatric speciation, where populations diverge in isolation due to limited gene flow across physical barriers (i.e., water). Divergence on isolated islands tends to follow an allopatric speciation model over evolutionary time (Rundell & Price 2009, Gillespie et al. 2020). Due to their geographic isolation and generally small size, island populations tend to lack zones of secondary contact between diverging avian taxa (Kinsey 1937, Diamond 1977), like those found in well-studied continental hybrid zones (Toews et al. 2016, Brelsford et al. 2017, Wang et al. 2020). Despite this generality, sufficient reproductive isolation and ecological divergence can evolve rapidly—even between sister taxa—in more isolated islands (e.g., Darwin’s finches, Lamichhaney et al. 2015). However, islands in the Indo-Pacific that are generally more species-rich and less isolated tend to produce instances of secondary sympatry between congeneric taxa that are attributed to colonization from other archipelagos, not from within archipelagos (i.e., intra-archipelago; Diamond 1977). As such, it is rare for island taxa to occur sympatrically with their closest relative (Mayr & Diamond 2001, Andersen et al. 2015, Cowles & Uy 2019, Manthey et al. 2020). Therefore, to better understand the challenges of intra-archipelago speciation, it is important to study taxa that come into secondary contact with previously isolated congeners.

Natural invasions between allopatric island lineages have rarely been witnessed. As such, there are few avian examples where secondary contact produces traditional hybrid zones on islands (Graves 2015, Sardell & Uy 2016). Instead, hybridization in island birds tends to be expressed as hybrid populations, an observation first made from careful study of museum specimens collected across Melanesia (Mayr 1932a, 1932b, 1938, 1942; Mayr & Diamond 2001). Recent genetic studies have provided further support for this phenomenon in islands and in other isolated areas of secondary contact (Nietlisbach et al. 2013, Lavretsky et al. 2015, Barrera-Guzmán et al. 2017, Andersen et al. 2021, Colella et al. 2021, McCullough et al. 2021). This is because islands lack the spatial scale to host a full hybrid zone and lack consistent parental input required for early generation hybrids (Kinsey 1937, Mayr 1942, Diamond 1977, Kissel and Barraclough 2010). Several aspects of island geography further facilitate the formation of hybrid populations. For example, the higher number of migrants originating from larger islands and continents can mean that smaller islands adjacent to larger ones are especially prone to the formation of hybrid populations (see MacArthur & Wilson 1963, 1967; Gyllenhaal et al. 2020). Additionally, glacial cycles can cause cyclical connectivity of islands and thereby increase rates of gene flow between island populations (Brown et al. 2013, Tan et al. 2022). This phenomenon of glacial eustasy could promote the formation of island hybrid zones, but as such cycles have occurred for millennia, present-day insular hybrid zones likely represent a deviation from longer-term isolation.

A conspicuous element of island bird faunas, especially in the Indo-Pacific, is the abundance of widespread ‘polytypic’ species. These geographic radiations occur on many islands—often across multiple archipelagos—and although apparently closely related, each island population may differ markedly in plumage pattern or coloration. *Pachycephala* whistlers are amongst the most widespread and strikingly diverse geographic radiation across the Indo-Pacific (Galbraith 1956) and Ernst Mayr leveraged this ‘great speciator’ lineage to inform our early understanding of the process of allopatric speciation (Mayr 1932a, 1942, 1963; Mayr & Provine 1980). Most insular populations of *Pachycephala* are allopatric, but there are several instances of secondary contact between divergent lineages, which has led to outcomes that range from secondary sympatry to apparent hybrid populations (Mayr 1932a, Mayr 1932b, Mayr & Diamond 2001).

The Fiji Whistler (*Pachycephala vitiensis*) is a monophyletic lineage of 10 named taxa (Andersen et al. 2014; Jønsson et al. 2014), which is part of the broader geographic radiation known as the Golden Whistler species complex (Galbreath 1956, 1967). Most taxa are single-island endemic populations that are distinct in appearance; yet, their distinctiveness consists of variations on a theme. All golden whistlers have dark backs and yellow underparts, but populations differ in combinations of throat color (white or yellow), the presence or absence of a black band across the breast, and other minor plumage details. Thus, three main types of populations exist: white-throated with a black breast band, yellow-throated with a black band, and yellow-throated with no band (Fig. 1). Typically, one such phenotype occurs across an archipelago (e.g., *P. citreogaster* in the Bismarck Archipelago or *P. chlorura* in Vanuatu); however, all three plumage types are represented within Fiji, and Mayr hypothesized that the Fiji Whistler represented ongoing contact between an old, yellow-throated lineage and a newly expanding, white-throated lineage. This hypothesis featured prominently in some of his influential works (e.g., Mayr 1942, Mayr 1963), especially how the failed secondary contact of this and other whistlers highlight the importance of reproductive isolation in permitting sympatry. Moreover, this lent support to his conclusion that allopatric lineages such as these do not represent biological species (Mayr 1932a). It was also used to support the notion that islands facilitate the formation of hybrid populations (Mayr 1942). Here, we use genomic data to test Mayr’s hypothesis of colonization history and assess the nature of admixture upon secondary contact between two phenotypically divergent lineages on Vanua Levu and its outlying islands in Fiji (Fig. 1A). We tackle these questions using three genomic datasets: restriction site-associated DNA sequencing (RAD-seq) of the focal Fijian taxa, target capture data of a broader range of taxa to determine the age of the Fijian radiation, and a new draft reference genome.

**Figure 1:**
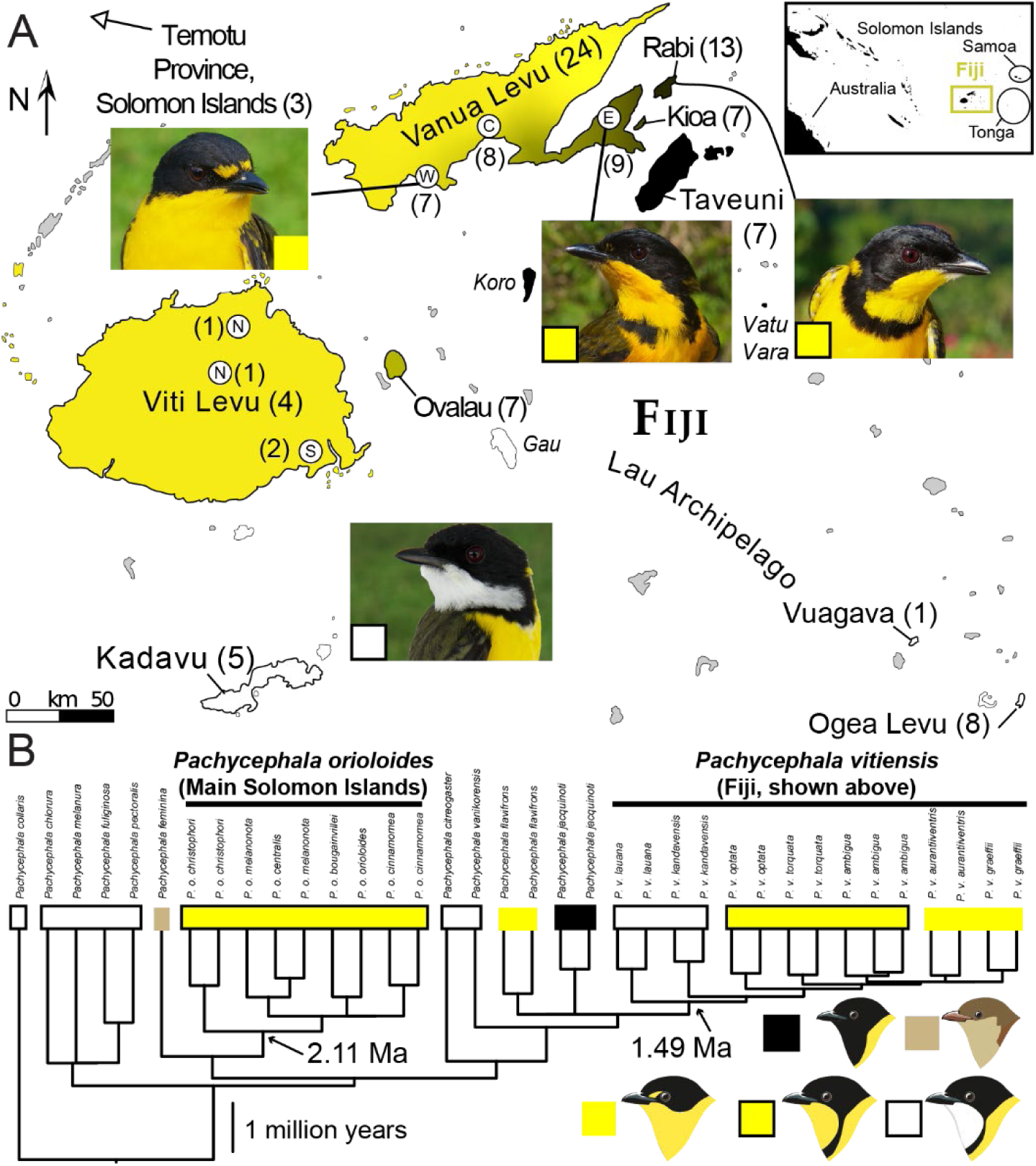
A) Map of islands sampled and a representation of plumage coloration. Islands with unknown plumage coloration are gray. Sampled islands are labeled by name and have a sample number. Unsampled islands with distinct taxa are labeled in italics. Islands without whistlers are gray, and island color corresponds with phenotype. White represents males with a white throat and black breast band, black represents a yellow throat and black breast band, yellow represents a yellow throat and no breast band, and intermediates between black and yellow represent rough frequencies of black breast band. Photos of archetypical individuals are shown, with a box corresponding with the color scheme in B. Sampling points (white circles) on large islands are shown, and when relevant the abbreviated cardinal direction is used to discuss samples from given points. Inset depicts the location of Fiji relative to other areas of interest. B) Phylogeny inferred from ultraconserved elements for more whistler species used for estimating divergence dates. Tips have boxes corresponding to male plumage patterns, with cartoon examples of phenotypes in the bottom right corner. Mean crown divergence times inferred by BEAST are at relevant nodes. Artwork by Jenna McCullough, some with modifications, based on artwork from Birds of the World (Billerman et al. 2020).

## Methods

### Sampling and DNA Extraction

We collected genetic data from all major phenotypes in the Fijian archipelago, with an emphasis on the contact zone across Vanua Levu (Fig. 1). We sampled 79 individuals of the genus *Pachycephala* for our primary RAD dataset (excluding samples with too few reads or too high relatedness), of which 76 comprised our core *P. vitiensis* sampling, and a non-mutually exclusive set of 63 samples comprised our yellow-throated ingroup sampling (Fig. 1). All individuals were represented by tissues with vouchered specimens (Table S1). For yellow-throated Fijian taxa (formerly *Pachycephala graeffii*), we sampled seven from Ovalau (*P. v. optata*), four from Viti Levu (*P. v. graeffii*), 24 from Vanua Levu (*P. v. aurantiiventris* and *ambigua*), 13 from Rabi (*P. v. ambigua*), seven from Kioa (*P. v. ambigua*), and seven from Taveuni (*P. v. torquata*). For white-throated taxa, we sampled five from Kadavu (*P. v. kandavensis*), eight from Ogea Levu (*P. v. lauana*), one from Vuagava (*P. v. lauana*), and three from Nendo (Solomon Islands; *P. vanikorensis ornata*). We lacked three single-island, subspecific populations of *P. vitiensis*: the yellow-throated *P. v. bella* and *P. v. koroana* from Vatu Vara and Koro, respectively, and the white-throated *P. v. vitiensis* from Gau. In addition to our primary RAD dataset, we used one sample of *P. v. kandavensis* (KU 117379) to generate a nuclear reference genome assembly using linked-read sequencing on the 10X chromium platform. Finally, we compiled a dataset of ultraconserved elements (UCEs; Faircloth et al. 2012) from 36 individuals across the *P. pectoralis* superspecies (Table S1). This UCE dataset represents an expansion of Clade A from Brady et al. (2022) and is composed of 24 newly sequenced samples and 12 samples sequenced for Brady et al. (2022). Four of these samples were derived from museum specimen toepads (two each of *P. jacquinoti* and *P. flavifrons*), while the rest were from specimen-vouchered tissues (Table S1).

### Reduced-representation Bioinformatics

We used Stacks v2.4.1 (Rochette et al. 2019) to call single nucleotide polymorphisms (SNPs) for our focal RADseq dataset and used the outputs as input for downstream analyses. We used the *process_radtags* module in single-end mode for demultiplexing reads by individual, which we then aligned to our reference genome using the *mem* algorithm of BWA v0.7.17 (Li & Durbin 2009). We used the Stacks module *gstacks* to assemble RAD loci based on these alignments and call SNPs. We generated multiple datasets using *gstacks* for different analyses and used the *gstacks* output to generate input files using *populations*. Unless otherwise stated, we performed these analyses with a 75% complete SNP matrix (i.e., SNPs present in 75% of individuals were included). We processed UCEs using the phyluce v1.7.0 pipeline (Faircloth 2015), following exact details as outlined in Brady et al. 2022.

### Population structure

We used several methods to explore population structure at all levels using RAD data, but with a focus on yellow-throated *P. vitiensis*. First, we calculated Weir and Cockerham’s (1984) F_ST_ between each sampling region with more than one individual (island or sub-region of island if relevant) using a custom wrapper around vcftools v0.1.15 (Danecek et al. 2011). Although Weir and Cockerham’s estimator can be biased slightly for low sample sizes at higher levels of divergence, this is not relevant for focal comparisons (Willing et al. 2012). This script was first generated for Mapel et al. (2021), and an updated version can be found on GitHub: https://github.com/ethangyllenhaal/FijiPachyRad. This script was run separately for loci assigned to autosomes and the Z chromosome. Second, we ran principal component analyses (PCAs) using the glPca module of adegenet v2.1.1 (Jombart 2008) in R v3.6.1 (R Core Team, 2019), which uses imputation to address missing data. Input was generated using the R package vcfR v1.8.0 (Knaus and Grünwald, 2017). Finally, we evaluated admixture proportions of yellow-throated *P. vitiensis* with sNMF (Frichot et al. 2014) in the R package LEA v2.2.0 (Frichot & François, 2015). Input for sNMF excluded uninformative singletons (i.e., minor allele count <2; Linck & Battey 2019). The alpha (normalization) parameter was set to 10 after exploratory runs of what minimized the cross-entropy criterion, but the results were similar across different values. The result with the minimum cross-entropy criterion after 50 iterations per k value was chosen for plotting.

### Phylogenetic trees and networks

We generated a phylogeny of our RAD data using a concatenated maximum likelihood approach (treating heterozygotes as equal likelihoods of either nucleotide) with IQ-TREE v2.8.3 (Minh et al. 2020), using ModelFinder (Kalyaanamoorthy et al. 2017) for model selection with 1000 ultrafast bootstraps (Hoang et al. 2018) for assessing support. We used individual-level, phylip-formatted data generated from a 90% complete matrix. Two samples presented as problematic (unexpected location based on other analyses, reducing support value of intervening samples) and were removed from subsequent sequence-based analyses. We also estimated a species tree using SVDQuartets in PAUP* v4.0a169 with 100 bootstraps and evaluating 100,000 quartets (Chifman & Kubatko 2014), grouping samples by sampling region (Fig. 1). To estimate the age of our focal radiation, divergence time dating was performed in BEAST v2.6.7 (Bouckaert et al., 2019; Appendix 1). To account for major introgression events, we generated a population tree with inferred admixture events using TreeMix v1.13 (Pickrell & Pritchard 2012). This analysis was conducted with a 90% complete matrix of SNPs to limit the effects of missing data. We iteratively added migration edges until the last meaningful edge was added that increased the likelihood by more than two (equivalent to a change of the Akaike Information Criterion of four, a common threshold for model selection).

### Tests for gene flow

We performed statistical tests for gene flow with ABBA/BABA tests as implemented in Dsuite v0.4r38 (Malinsky et al. 2021) with *P. vanikorensis* as an outgroup, with a 90% complete VCF as input. We calculated the D statistic (Durand et al. 2011) and its significance in addition to the f_4_ estimate of admixture proportion (Patterson et al. 2012). We focused on two *a priori* hypotheses: 1) gene flow between populations on Vanua Levu and those on nearby islands and 2) gene flow between white- and yellow-throated *P. vitiensis*. We also used the f_branch_ function to make inferences about timing of gene flow (Malinski et al. 2018). We took three approaches to account for multiple comparisons, described in Appendix 1 (Supplemental Methods; Correcting for multiple tests for gene flow).

### Genome scans

To understand the genomic architecture of phenotypic divergence, we estimated Weir and Cockerham’s (1984) weighted F_ST_ in 10 kbp windows across the genome using vcftools v0.1.15 (Danecek et al. 2011) and plotted the results using the R package qqman v0.1.8 (Turner 2018). In most cases (but not all), this windowed value is the mean from one RAD locus; windows without SNPs are ignored. We took two approaches to assign comparisons: geographic locality and phenotype. For the former, this represents a standard inter-taxon F_ST_ outlier comparison, and the latter is closer to a GWAS. The phenotypic indices were estimated by MJA, and included indices for the extent of yellow in the lores and black on the breast of yellow-throated *P. vitiensis*. In short, the extent of the band was binned into four categories (absent, hooked, dirty, and complete; 0–3 respectively) and lores into three (black, minimally yellow, and completely yellow; 0–2 respectively). We assessed putative functional genes based on our annotations (Supplemental Methods) for windows with F_ST_ greater than 0.5 in the eastern vs western Vanua Levu, comparison, including a region bounded by values >0.5 on the Z chromosome.

## Results

### Population structure

Each of the three white-throated *P. vitiensis* populations formed distinct genomic clusters, all of which were notably distinct from the yellow-throated populations (Fig. S1). Within the yellow-throated taxa, there was notable population structure corresponding to geography and the extent of dark plumage in the breast and lores of a given population (Fig. 2C– D, S3). In principal component space, more positive values of PC1 largely corresponded to darker populations. The one exception was the population on Ovalau, which has intermediate plumage but was separated in PC space from all other yellow-throated populations (Fig. 2C–D). Populations from Kioa and eastern Vanua Levu clustered together, as did populations from central and western Vanua Levu. Estimates of admixture fractions tended to be primarily from only one ancestral population, but notable levels of mixed ancestry were consistently inferred for eastern Vanua Levu (Fig. 2D, S3). Estimates of genome-wide F_ST_ across Vanua Levu were associated with plumage coloration. For example, with the western Vanua Levu populations as a reference, F_ST_ increased moving east across Vanua Levu, with a further increase on the darker island populations on Kioa, Rabi, and Taveuni (Table S3). F_ST_ was generally higher on Z-chromosomes than on autosomes, except for the comparison between Kioa and eastern Vanua Levu.

**Figure 2:**
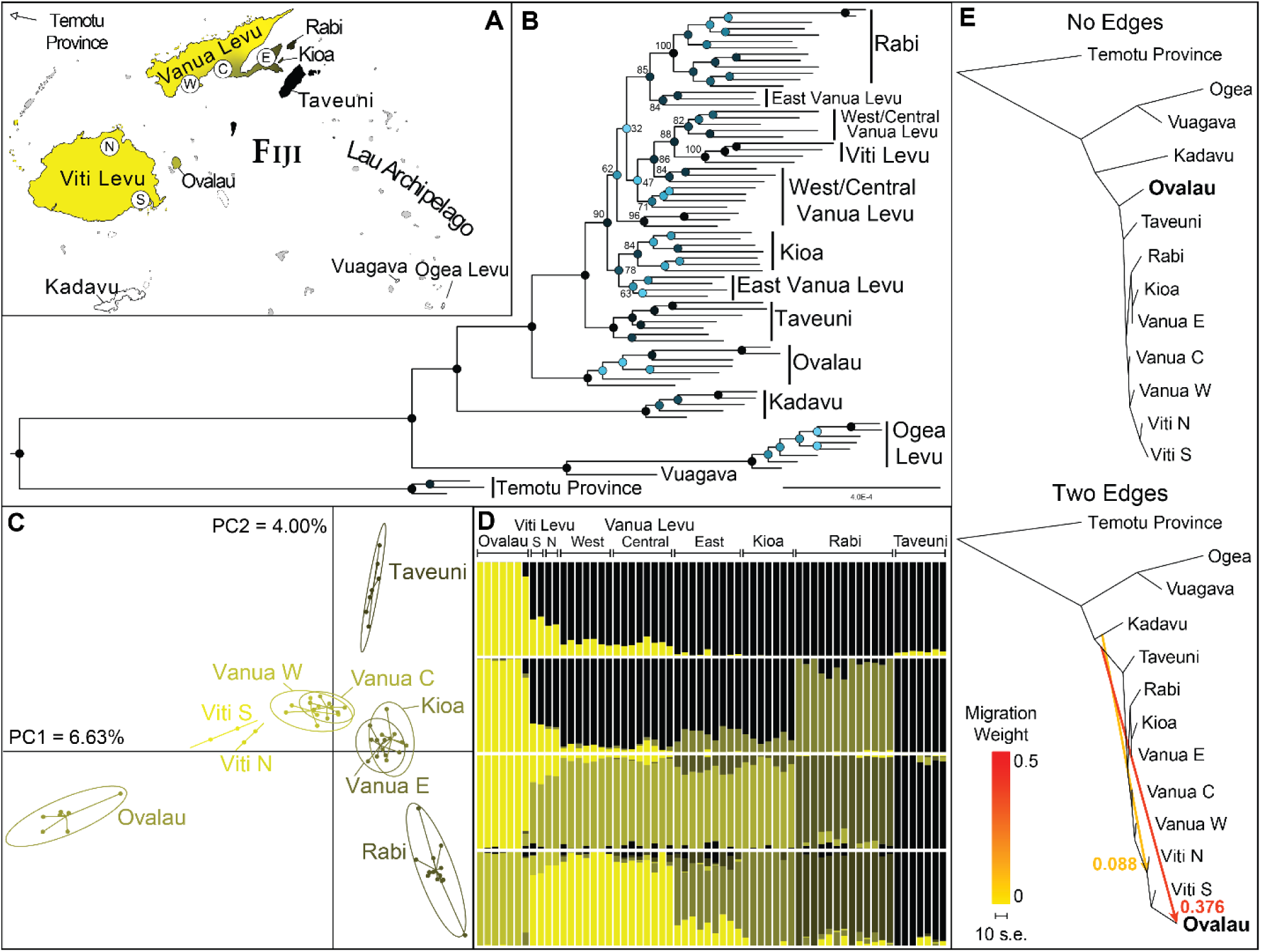
A) Map of focal sampling localities in Fiji. North, South, East, West, and Central are abbreviated to N, S, E, W, and C, respectively. B) IQ-TREE Phylogeny of RAD-seq dataset, with groups of tips labeled corresponding to the inset map. Node shapes correspond to support value, with lighter values indicating lower support. Support values for potentially geographically informative nodes in the main yellow-throated clade are shown, but we note relationships among the main yellow-throated clade are largely unresolved. C) Genomic PCA of all yellow-throated taxa, with order of colors on a yellow-black gradient corresponding to the relative mean band extent in the population (more extensive bands in blacker clusters). D) Ancestry plots from sNMF analyses of yellow-throated taxa for K values of 2-5, colored as C. E) TreeMix Phylogenetic networks with zero and two migration edges, with labels corresponding to the weight of the edge (see Figure S3 for one edge network).

### Phylogenetic trees and networks

Our phylogeny inferred in IQ-TREE recovered the white-throated Lau populations as sister to other *P. vitiensis*, with the population on Kadavu sister to all yellow-throated taxa (Fig. 2B, Figure S2). Within the yellow-throated populations, Ovalau individuals were sister to all others, then Taveuni was moderately (bs= 90) supported as sister to the remaining yellow-throated clade. Support values for the rest of the tree were low, and many populations that clustered in PCA space were not monophyletic. Populations that were fully supported as monophyletic were from the relatively isolated islands of Rabi and Viti Levu. The phylogenetic network from TreeMix recovered a similar topology when no migration edges were added (Fig. 2E, S3). However, when migration was modeled, a significant migration edge was inferred between Ovalau and an ancestral taxon (admixture fraction=0.376; p < 0.0001). In turn, Ovalau’s placement in the tree changed to be sister to the southern Viti Levu population and nested within the clade of populations with weaker breast bands (Fig. 2E). A second, less significant (p = 0.0067), edge was inferred between an ancestral Kadavu population and the ancestor of the Viti Levu clade when two edges were considered (Fig. 2E). Our species tree analysis in SVDQuartets produced some similar results but was very sensitive to the inclusion of the Ovalau population (Fig. S4). Although several nodes were decently supported without it (often more so than in IQ-TREE), the putatively admixed population resulted in high topological uncertainty when it was included. Our UCE time tree (Fig. 1B) suggested that the crown age of Fijian whistlers is approximately 0.75–2.58 Ma (mean 1.49 Ma) and that mean ages of island-specific taxa in Fiji range from 0.66–0.96 Ma. Within the Solomons Whistler (*P. orioloides*), there was general support for clades corresponding to geologically cohesive archipelagos within the broader Solomons Archipelago—the New Georgia Group, the Bukida Group, and Makira Island (excluding *P. feminina*)—which had an inferred crown age of 0.89–3.28 Ma (mean 2.11 Ma). The UCE topology was similar for the Fijian clade despite lower sampling (Fig. S5).

### Tests for gene flow

Tests for gene flow in the group provided evidence for widespread gene flow throughout the archipelago, although comparisons within closely related groups were not possible due to the constraints of ABBA/BABA tests. The significant tests (p < 0.05) between adjacent populations are outlined in Figure 3A. All tests are in Table S3, and f_branch_ in Figure S6. In short, high levels of gene flow (f_4_ admixture fraction > 0.2) were inferred between Kioa and western and central Vanua Levu, Rabi and points west, Taveuni and both Rabi and Kioa, and Ovalau and Viti Levu. The latter was highly significant, likely due to the admixed nature of Ovalau, and resulted in unusual topologies being tested. All of these were significant after accounting for multiple comparisons except for gene flow between Rabi and points west (harmonic p=0.068) and between eastern Vanua Levu and the rest of Vanua Levu (harmonic p=0.053). However, the latter is supported by phenotypic evidence. Low levels of gene flow were inferred between Viti Levu and Kadavu and between Ogea Levu and several other populations (Viti Levu, central Vanua Levu, Rabi, and Kioa). These results are not robust to multiple comparisons, but gene flow between Viti Levu and Kadavu (or a related population) is supported by TreeMix (Fig. 2E).

**Figure 3.**
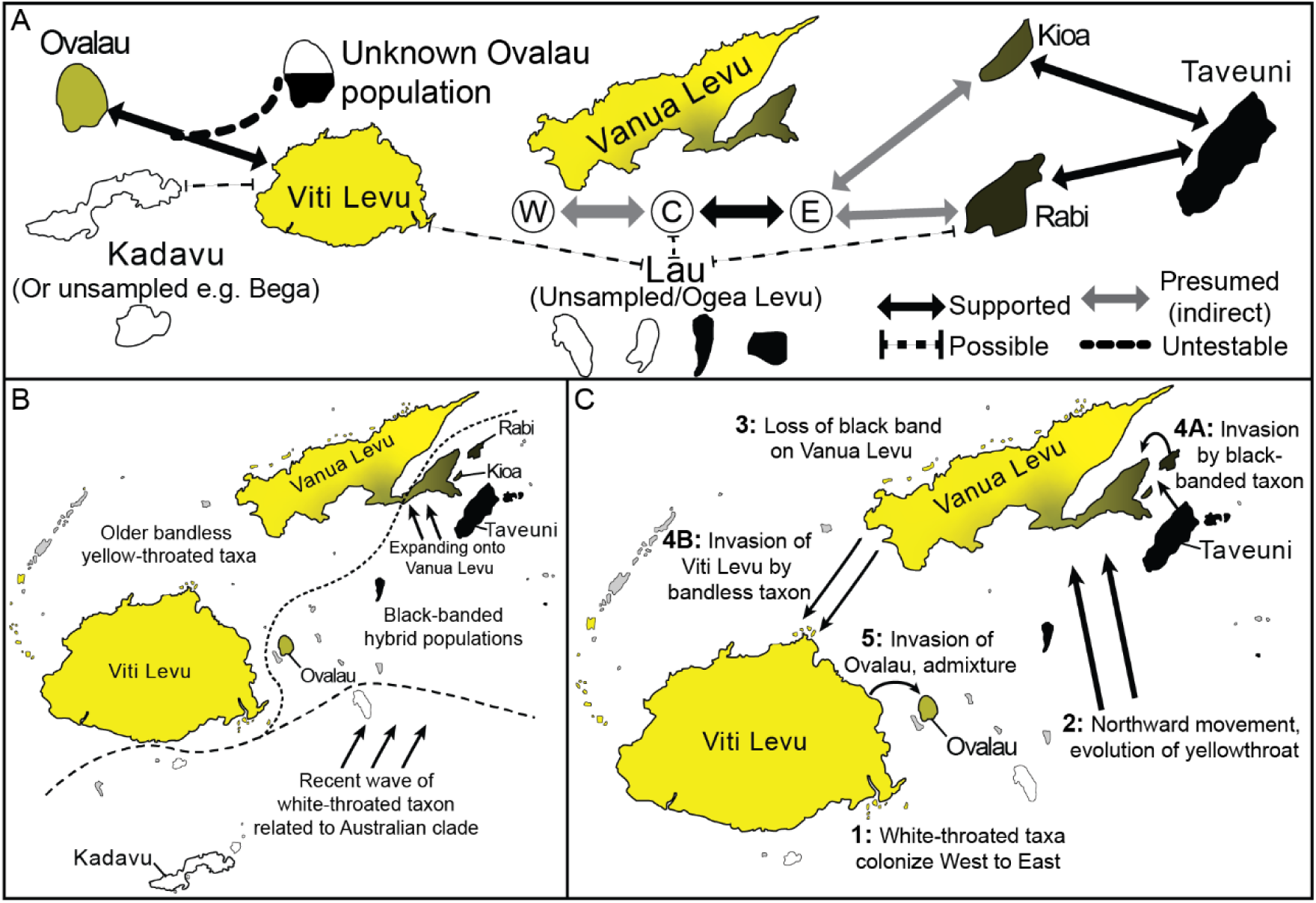
A) Sampled populations (represented by their respect islands’ shape and phenotype) with arrows corresponding to admixture fraction as estimated by the f4 ratio. The “possible” line denotes edges with significant gene flow for some tests but did not stand up to multiple comparisons. “Presumed” edges represent cases where gene flow is readily testable, but would have occurred based on gene flow inferred between non-adjacent populations (e.g., between Rabi and Western Vanua Levu). Note that the islands are not drawn to scale and are arranged to illustrate the linear pattern of inter-island gene flow. B-C) Diagrams of Mayr’s (B) and our (C) hypothesized colonization history of whistlers in Fiji.

### Genome scans

Our exploration of the genomic architecture of divergence found one consistent (multiple consecutive windows with high F_ST_ in the eastern vs western Vanua Levu comparison) peak differentiating darker and yellower populations, based on both geography and hybrid index. This peak resided on the Z chromosome and was clearest between the populations with the most similar genome-wide F_ST_ but differing band extracts (eastern vs western Vanua Levu; Figure 4). Several other peaks between Taveuni and other populations were also found, including a broader peak on the Z chromosome (Fig. S7). Our assessment of potential causal loci found two genes that have been correlated in plumage or pigmentation in past studies. One gene, TRPM7, is in the same family as TRPM1, which has been correlated with feather pigmentation in domesticated chickens. More notably, PCSK5 has been directly correlated with variation in melanin levels in wild birds, and is associated with melanin production pathways (San-Jose et al. 2017). However, several genes associated with other phenotypes such as fat deposition (NR2F2; Zhu et al. 2021) and stress responses (MCTP1; Taff et al. 2019) were also in this large region.

**Figure 4.**
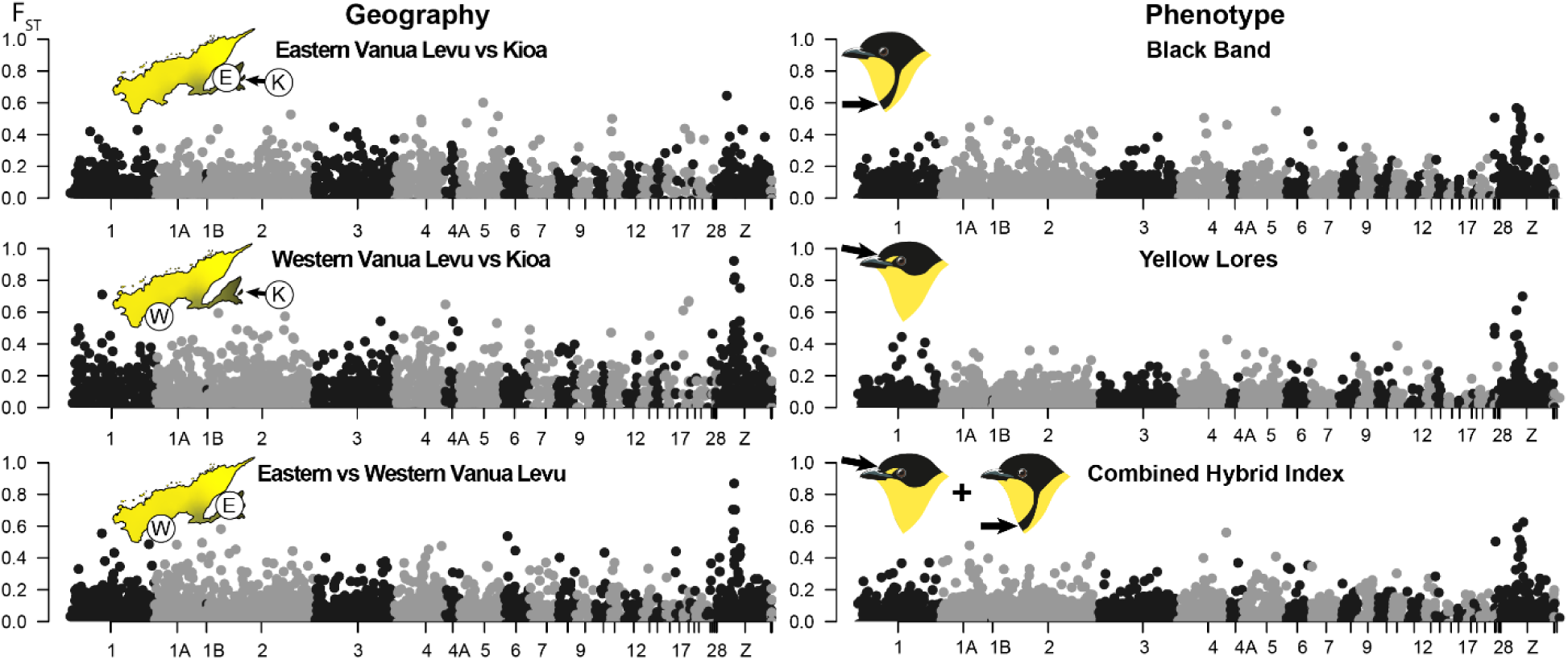
Manhattan plot of FST across the whole genome (based on synteny mapping with Zebra Finch). The left column compares by population (with East Vanua Levu and Kioa sharing similar phenotypes), while the right compares by phenotype. For the right column, the values compares are band values of 0–1 versus 2–3, lore values of 0 versus 1–2, and combined index values of 0–1 versus 3–5 (see Methods; Genome Scans).

## Discussion

The genus *Pachycephala*, particularly those in Fiji, had a major impact on the development of Ernst Mayr’s view of allopatric speciation during the Modern Synthesis (Mayr 1932a, Mayr 1932b, 1942). He interpreted phenotypic evidence to suggest multiple instances of hybrid populations after phenotypic divergence in allopatry, representing the failure of such lineages to attain reproductive isolation despite having plumage traits as different as some continental sympatric species in the same genus (e.g., *P. soror* and *P. schlegelii*, *P. pectoralis* and *P. melanura*). Our study supports the general idea that these birds failed to speciate completely when in allopatry, but we lacked support for Mayr’s exact hypothesis for how this occurred geographically (Fig. 3B). We instead present a hypothesis that emphasizes rapid plumage evolution occurring in geographic allopatry as the species colonized Fiji, producing much greater diversity in male plumage than seen in the older radiation in the Solomon Islands (Fig. 3C). However, a second wave of movement—likely facilitated by glacial cycles—resulted in secondary contact, which caused the formation of a hybrid population on Ovalau (Fig. 2, 3) and a series of intermediate populations across Vanua Levu (Fig. 2). We also find evidence that the phenotypic divergence in the Vanua Levu populations is largely tied to one region of the Z chromosome, rather than at several points across the genome.

### Secondary contact of allospecies

Our dataset included two instances of secondary contact, the contact zone on Vanua Levu and a genomically confirmed hybrid population on Ovalau, both recognized by Mayr as potential instances of contact between allospecies (Mayr 1932a). The Vanua Levu contact zone is between the two yellow-throated phenotypes, where a black-banded population invaded un-banded populations from outlying islands to the East (i.e., Kioa and Rabi), with high direct or indirect gene flow inferred between all sampled populations in the region (Fig. 4a). We did not find a clean clinal pattern, but rather a divide in phenotype and genotype between the two western Vanua Levu populations and eastern Vanua Levu plus Kioa. This suggests that further sampling of the southern coast of Vanua Levu is required to fully understand the hybridization dynamics in this group, but our detection of high levels of gene flow and low genetic divergence across the island suggests high admixture (Fig. 2, 3a; Table S3). The narrow stretch of land between the eastern and central sampled populations may have caused this relatively steep transition, and indeed may facilitate the long term persistence of a hybrid zone, as is seen in some continental contact zones (e.g., Cicero et al. 2023).

For Ovalau, a population noted by Mayr to harbor extensive phenotypic variability (Mayr 1932a), we find evidence for a lineage merger between an un-banded, yellow-throated population from Viti Levu and an unsampled or ghost taxon that likely originally occupied the island prior to this merger (Fig. 3, 4). Traditional concatenated and species tree analyses struggled with this taxon, either placing it as sister to other yellow-throated taxa with full support (Fig. 2) or resulting in inflated topological uncertainty (Fig. S4), respectively. TreeMix, which allows the inference of admixture events, resulted in a major topological rearrangement. Andersen et al. (2021; described further in McCullough et al. 2021) found a similar result in a strikingly similar geographic context, an isolated island fully or nearly connected during glacial periods, which along with other findings could suggest this is a frequency pattern in island taxa (Nietlisbach et al. 2013, Lavretsky et al. 2015, Colella et al. 2021). Much like the example in Andersen et al. (2021), the phenotype of the original Ovalau lineage is not fully known. It could either be a yellow-throated, black-banded population like those found on the unsampled island of Koro, or a white-throated, black-banded population like those found on Beqa and Gau islands. A better-documented hybrid population between yellow-throated and white-throated taxa has many individuals with intermediate throat color unlike the pure yellow throat of Ovalau individuals (Mayr 1932b). Further sampling should reveal whether this represents a population akin to Mayr’s hypothesis for the genetic origin of all black-banded birds, where hybridization between un-banded and white-throated populations produces the yellow-throated, black-banded phenotype that we sampled from islands east of Vanua Levu (Fig. 1A; Mayr 1932a, 1942). However, we found no evidence that the majority of yellow-throated, black-banded taxa represent hybrid populations (Fig. 3).

### Genomic architecture of allospecies divergence

We detected only a single multi-locus divergent genomic region in the instance of secondary contact on Vanua Levu (Fig. 4). This Z-linked region stood out the most strongly when only geography (i.e., western Vanua Levu compared to Kioa) was considered, but was evident when phenotype was considered as well. Although one gene in this region (PCSK5) has mechanistic ties to plumage divergence (San-Jose et al. 2017), it is part of a larger region full of genes involved in other pathways. Although it is tempting to ascribe causality to this gene, we believe further validation is required to rule out hypotheses such as genetic hitchhiking. Indeed, even if it is causal, the phenotype itself may not have evolved under strong selection, but due to alternatives such as hitchhiking off selection on a linked gene or drift (Gould & Lewinton 1979, Barton 2000). Regardless, inter-population divergence being linked to a single chromosomal region is consistent with genome-wide admixture proportions matching geography rather than within-population phenotype (Fig. 2C). Additionally, when the darkest birds on Taveuni were compared to other populations, including the relatively dark individuals on Rabi, a wider peak in the same chromosomal region was observed (Fig. S7). Future work can assess if reduced recombination across lineages drove divergence in this region, adaptive or not. The low number of genomic regions involved in divergence is also consistent with the apparent lack of resistance to gene flow, as even if selective forces shape this divergence, polygenic adaptation better promotes cohesion during secondary contact (Barton 1983, Barton & Hewitt 1985, Flaxman et al. 2014). If this region is broadly responsible for the similar plumage diversity in other whistlers, it may suggest a model where great speciators evolve high plumage divergence despite minimal intrinsic reproductive isolation or ecological specialization.

### History of colonization and plumage divergence

Our results provide support for two themes of Mayr’s hypothesis: dynamic waves of colonization and the lack of reproductive isolation between allopatric lineages (Mayr 1932a, 1942). However, the underlying dynamics of colonization that we inferred differed starkly from Mayr’s interpretation. Mayr hypothesized that recent expansion of white-throated taxa into the Fijian archipelago from Vanuatu (*P. chlorura*), resulting in the introgression of a dark-banded phenotype into an unbanded, yellow-throated population and the formation of hybrid populations of banded, yellow-throated birds (Mayr 1932a). We found no evidence of recent contact between “old” and “new” phenotypes. This is evidenced by the monophyly of yellow-throated birds and the nested nature of less banded birds in our phylogeny (Fig. 3). Indeed, the proposed white-throated lineage does not represent a monophyletic group, but rather a laddered series of white-throated taxa leading to the yellow-throated clade. We also found no evidence that banded, yellow-throated populations, specifically those on Taveuni, shared an excess of alleles with white-throated lineages (Fig. 3C, 4A, S4).

We propose an alternative model for the colonization of Fiji whistler. First, a white-throated lineage underwent an initial wave of colonization of the Fijian archipelago, followed by the evolution of both yellow-throated phenotypes and a second wave of dispersal among entirely (or mostly) yellow-throated populations, which led to the two admixture scenarios outlined above (Fig. 4C). The phylogenetically nested nature of yellow-throated Fijian populations within a clade of white-throated Kadavu and Lau populations (Fig. 1A, 3B) suggests a wave of colonization by a dispersive, initially white-throated lineage throughout isolated islands of Fiji. Second, the loss of a white throat from northern and western Fijian populations suggests a single evolution of a yellow-throated phenotype during the northward colonization of the archipelago. Third, the nested nature of unbanded populations suggests the loss of the black band while outlying islands (e.g., Koro, Taveuni, and Rabi) retained their banded phenotype. Fourth, we propose a recent wave of admixture driven by glacial cycles resulting in increased propagule pressure from Rabi and Kioa to the Natewa Peninsula of Vanua Levu—which is relatively isolated from the rest of the island due to the long, narrow base of the peninsula—resulting in a zone of phenotypic intermediates. The glacial cycles then also resulted in the colonization of Viti Levu by this clean-breasted form, possibly filling an empty niche or displacing and genetically swamping a white-throated ghost population related to birds on Kadavu (resulting in a signal of gene flow, Figure 3C and 4A). This last point is supported by the close relationship between Viti Levu and western Vanua Levu populations, despite the long over-water distance during interglacial periods, and continued separation during glacial periods (Fig. 2, Table S2). Finally, during this wave or during glacial maxima, the island of Ovalau, was partially genetically swamped by the new clean-breasted lineage that had colonized Viti Levu. We believe that the relatively distinct nature of Ovalau, an island that was fully connected to Viti Levu during glacial maxima, is consistent with this admixture occurring during a recent glacial cycle, as the propagule pressure from extended contact would likely fully erode the genetic and phenotypic signatures of the population’s admixed origin. Indeed, once gene flow is accounted for, Ovalau is nested within the Viti Levu populations (Fig. 3C). Estimating the timing of these events in a phylogenetic context is complicated by intraspecific divergence (Fig. 1B) and confounding gene flow (Fig. 4). However, our proposed historical model is supported by multiple lines of genetic evidence and is parsimonious with respect to the number of colonization waves and amount of gene flow between extant taxa.

This colonization pattern is unlike another taxon with a similar distribution, the wattled honeyeaters (*Foulehaio*). In this case, older lineages occupy the larger islands of Fiji, while more isolated islands in the Lau, Tongan, and Samoan archipelagos are recently diverged with evidence of long-distance gene flow between them (Mapel et al. 2021). Instead, *Pachycephala* illustrates the opposite pattern, with populations in Lau, Tonga, and Samoa sister to the populations on the larger islands (Fig. 1b, 2). However, the genetic and phenotypic affiliation between the Natewa Peninsula and Taveuni is consistent with the endemism of *Lamprolia* silktails (Aves: Rhipiduridae)—a deeply divergent relictual lineage—to those two regions (Irestedt et al. 2008, Andersen et al. 2017). These observations highlight that although we can make some predictions about how recently diverged, allopatric populations will interact upon secondary contact, colonization history can be highly idiosyncratic between co-distributed lineages.

## Conclusion

Our work revisited a system that had a disproportionately large influence on Mayr’s contributions to the Modern Synthesis and the study of speciation (Mayr 1942). Fiji whistlers feature many important elements of speciation theory. Most relevant to Mayr’s implementation on the Biological Species Concept, we found a lack of evidence for reproductive isolation of allopatric taxa despite having phenotypic divergence comparable to sympatric species (Mayr 1942). Relating to the study of the genomic architecture of speciation, we found one instance of secondary contact that resulted in high introgression across Vanua Levu (Fig. 4A) demonstrated the weakness of single genomic regions in producing strong reproductive isolation (Barton 1983, Flaxman et al. 2014). Finally, as investigators have increasingly demonstrated in recent decades, geographic variation in plumage coloration can evolve rapidly, with two major transitions in plumage phenotype occurring in a short span of time (approximately 500,000 years). This is especially impressive compared to the *P. orioloides* radiation in the Solomon Islands, which has mostly undergone changes in the extent of black plumage over a longer period of time than it took for *P. vitiensis* to evolve its wide range of phenotypes (Fig. 1B). Our study posits a new biogeographic hypothesis for the Fiji whistler and establishes a framework to inform future work on the genomic basis of color evolution in this group. Finally, our study highlights the value of historic and modern collections to reexamine long-standing hypotheses about how species form and diversify.

## Supporting information

RADSeq sampling table

UCE sampling table

## Acknowledgements

We are indebted to the staff and curators in the South Pacific Regional Herbarium at the University of the South Pacific, Suva (Marika Tuiwawa and Tokasaya Cakacaka), the Fiji Department of Forestry (Sanivalati Vido), the Biosecurity Authority of Fiji (Joeli Vakabua), Mika Bolakania, and Dick Watling for their assistance, permission, and friendship in Fiji. We are grateful to the following people and institutions for loaning new samples necessary for this project: the Mark Robbins (University of Kansas Biodiversity Institute); Paul Sweet (American Museum of Natural History); and Sharon Birks (University of Washington Burke Museum, USA). We are also grateful to the following people for loaning samples originally sequenced in Brady et al. 2022: Jack Dumbacher and Maureen (Moe) Flannery (California Academy of Science); Alex Drew (Australian National Wildlife Collection), and Robb Brumfield (Museum of Natural Science at Louisiana State University). We thank Jenna McCullough, Nicholas Vinciguerra, David J.X. Tan, and Elizabeth Solis, whose comments helped improve the manuscript. We would like to thank the UNM Center for Advanced Research Computing, supported in part by the National Science Foundation, for providing the high performance computing resources used in this work. We thank the KU Genome Sequencing Core (supported by National Institutes of Health grant 5P20GM103638 to E.A. Lundquist) for access to lab equipment and services. We gratefully acknowledge funding from the NSF’s Graduate Research Fellowship (DGE-1650114) and NSF awards to MJA (DEB-1557051, DEB-2112467) and RGM (DEB-1557053).

## Data Accessibility Statement

Population genetic input files, phylogenetic trees and input files, and reference genomes will be uploaded to Dryad: https://doi.org/10.5061/dryad.k98sf7mft. Scripts are archived on Zenodo, linked to the Dryad repository, as well as on a personal GitHub: https://github.com/ethangyllenhaal/FijiPachyRad. Raw Illumina sequencing reads for RAD-seq and new UCE data are available from the NCBI SRA (BioProject PRJNA1088558).

