## Appendix for "Waves of Colonization and Gene Flow in a Great Speciator"

### Repeated Waves of Colonization of a Great Speciator

#### Supplemental Methods

##### *DNA extraction, library prep, and sequencing*

For the focal RADseq dataset, we extracted genomic DNA using QIAgen DNeasy extraction kits (Qiagen Inc. Valencia, CA, USA) following the manufacturer's protocols. We quantified DNA with a Qubit 3.0 Fluorometer and visualized the quality of each sample using agarose gel electrophoresis. Whole genomic DNA extracts were standardized to 5 ng/µL and plated at 10 uL per sample (50 ng of DNA). Multiplex shotgun genotyping RADseq libraries (Andolfatto et al. 2011) were prepared at the KU Genome Sequencing Core facility. Briefly, genomic DNA was digested using the enzyme NdeI, bar-coded adapters were ligated to samples to allow for multiplexing, DNA fragments were size-selected in the 495–605 bp range, size-selected fragments were amplified using PCR, and amplified samples were cleaned using AMPure beads (Beckman Coulter) to remove primer dimers and short-fragment DNA. Single-read 100 bp Illumina sequencing was performed on a High Output run of a NextSeq 550 machine. UCE data was generated following the protocol in Brady et al. 2022, which, in turn, followed methods outlined in Faircloth et al. 2012. In short, Illumina libraries were prepared using a Kapa Biosystems Hyper Prep kit and then enriched for UCEs with a MYcroarray MYbaits probe kit.

##### *Reference Assembly*

High molecular weight genomic DNA was extracted from muscle tissue at HudsonAlpha Institute for Biotechnology and prepared into a 10X Chromium sequencing library. This library was sequenced with 150 base pair (bp) paired-end reads across two lanes on an Illumina HiSeq X machine, which generated 337,546,830 linked read pairs passing initial quality filters. We used the Supernova *de novo* genome assembler (Weisenfeld et al. 2017) with default parameters to assemble these linked 10X Chromium reads into preliminary scaffolds and to resolve phased haplotypes. Because of the conservation of genomic architecture among avian lineages (Ellegren, 2010) and the contiguity of our *Pachycephala* genome assembly, we next used syntenic mapping to assign our Supernova scaffolds to putative chromosomal positions. We ran Satsuma v1 (Grabherr et al., 2010; https://satsuma.sourceforge.net) using one of the haplotype-resolved assemblies output by Supernova and the *Taeniopygia guttata* reference genome assembly (version: TaeGut3.2.4, available from: http://ftp.ensembl.org/pub/release-97/fasta/taeniopygia_guttata/dna/) as input with default parameter settings. Satsuma performs genome-to-genome alignment of a fragmented query genome against a chromosome-level reference assembly using a combination of cross-correlation with fast Fourier transform, a match-scoring scheme that is robust to false hits, and an asynchronous 'battleship'-like search that permits computationally efficient genome comparisons. The resulting scaffolded query genome is represented by superscaffolds and pseudochromosomes which comprise contiguous sections of sequence relative to the reference genome sequence. This synteny mapped assembly was used here for aligning our single-digest restriction site associated DNA (RAD) loci.

##### *Reference annotation*

To assess potential functional genes underlying intra-subspecific divergence, we annotated our new reference genomes, taking inspiration from the pipeline used in Eliason et al. 2022. We used our pseudochromosome-level genome as input for RepeatMasker v4.1.5 (Smit et al. 2015), using repeats modeled after Zebra Finch (*Taeniopygia guttata*). Next, we performed structural annotation using a homology-based approach implemented in GeMoMa v1.9 (Keilwagen et al. 2018). We used two high-quality passerine genome annotations for inference: Zebra finch (*Taeniopygia guttata*; Warren et al. 2010) and Hooded crow (*Corvus cornix*; Poelstra et al. 2014).

##### *Divergence dating*

We used BEAST v2.6.7 (Bouckaert et al., 2019) to infer a time-calibrated tree. We used a birth-death tree prior, a lognormal, uncorrelated relaxed clock model, and assigned the HKY+G model of sequence evolution to each UCE locus. To reduce computing time, we constrained the topology to match our RAD tree using 14 most recent common ancestor (MRCA) priors in BEAUti v2.6.7. We parsed the 95%-complete data matrix into 10 subsets of 92 loci chosen at random without replacement, following McCullough et al., 2019 and Brady et al., 2022. We ran two independent Markov chain Monte Carlo (MCMC) chains of 50 million generations for each of 10 subsets, sampling trees every 10,000 generations. We visualized traces of log files in TRACER v1.7.1 (Rambaut et al., 2018) to assess convergence of individual runs and we ensured ESS values of all sampled parameters exceeded 200. We used LogCombiner v2.6.7 to combine two replicates from each of the 10 datasets and discarded the first 25% of trees as burn-in (50% burn-in for run 4). To produce a single maximum clade credibility (MCC) tree, we used TreeAnnotator v2.6.7 (Bouckaert et al., 2019) and subsampled every third tree for a final posterior distribution of 30,394 total trees.

There are no known fossil whistlers, so we relied on alternative calibration methods. We calibrated the two oldest nodes by establishing normal distributions using mean node ages derived from Brady et al., 2022, which were in turn, based on two secondary fossil-calibration points from Oliveros et al. (2019). Oliveros et al. (2019) used 13 fossil calibrations to date their passerine time tree, of which the closest fossil to Pachycephalidae was *Kurrartapu johnnguyeni*, a crown artamid that shared a most recent common ancestor with whistlers about 22.9 Ma. The first calibration point we used was for the crown group of all 36 tips, which we set with a normal distribution, sigma value of 0.5, and a mean date of 5.04 Ma (CI = 4.06–6.02). The second point was the split between *P. collaris* and remaining ingroup taxa with a normal distribution, sigma of 0.4, and a mean date of 3.48 Ma (CI = 2.7–4.26).

##### *Correcting for multiple tests for gene flow*

To confirm the significance of a given class of edge (e.g., between Kioa and points west, or Kadavu and all yellow-throated birds), we performed multiple methods that account for multiple comparisons. First was a heuristic method, based on the consistency recovery of significant tests for trios where the sister pair (P1 and P2) have notable population structure and P3 is geographically adjacent. We also calculated a harmonic mean p-value for a given edge class. We focus on this value for statistical accounting for multiple comparisons, but also corrected for comparisons within a class of edge using the Benjamini-Hochberg approach. These tests were performed after the removal of an admixed population (Ovalau), as its placement inflated significance of many tests.

#### Supplementary Results

##### *Sequencing information and RAD-seq bioinformatics*

Our sampled individuals had a mean per-sample coverage of 108.1x (range of 24.8–361.9x). The mean number of sites per locus for our full dataset was 70. After filtering, our full dataset resulted in 16,082 loci with 1,150,004 sites (i.e., approximately 1% of the genome), including 12,081 variable sites. Other datasets with different subsets and completeness ranged from 13,494–16,503 loci (including invariant loci) and 7,724–11,293 variants depending on the taxa included (Table S1) and completeness. This does not include the highly variable number of variants used for pairwise $F_{ST}$ estimation. Our 95% complete UCE matrix had 922 loci and 1,069,836 bp; mean locus length was 1,160.3 bp [range: 334–1,581 bp].

##### *Correcting for multiple tests for gene flow*

In correcting for multiple comparisons, we found that our primary conclusions are robust, but introduce caveats for certain cases where gene flow is not ongoing. All metrics of multiple comparisons supported gene flow from Taveuni to populations to the west (namely Rabi and Kioa) and gene flow from Kioa into central and western Vanua Levu populations. Gene flow between Rabi and Vanua Levu and eastern and other areas of Vanua Levu were only supported by the heuristic approach (although were marginally significant with harmonic p=0.068 and 0.053, respectively). Gene flow from eastern to other areas of Vanua Levu was likely not significant due to low power of the test, as it is clearly supported by phenotypic evidence and other tests. Most importantly, gene flow between white-throated (i.e., Lau or Kadavu) and yellow-throated populations was not robust to multiple comparisons.

Supplemental Figures and Tables

Figure S1: Principal Component Analysis (PCA) of all samples, with all yellow-throated *Pachycephala vitiensis* displayed as one group.

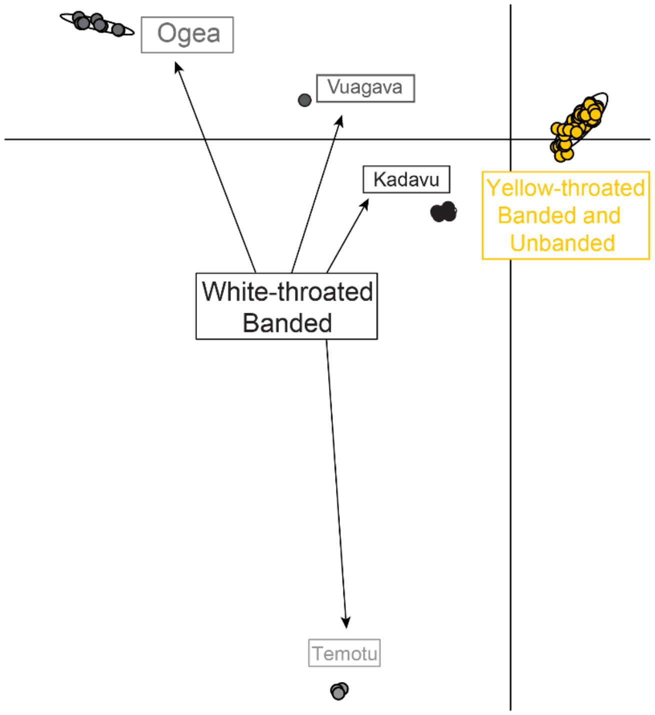


Figure S2: Full IQTREE analysis with all bootstrap support values (on nodes) and sample names (on tips). Sample names are abbreviated (Pv = P_vitiensis, Pg = P_graeffii).

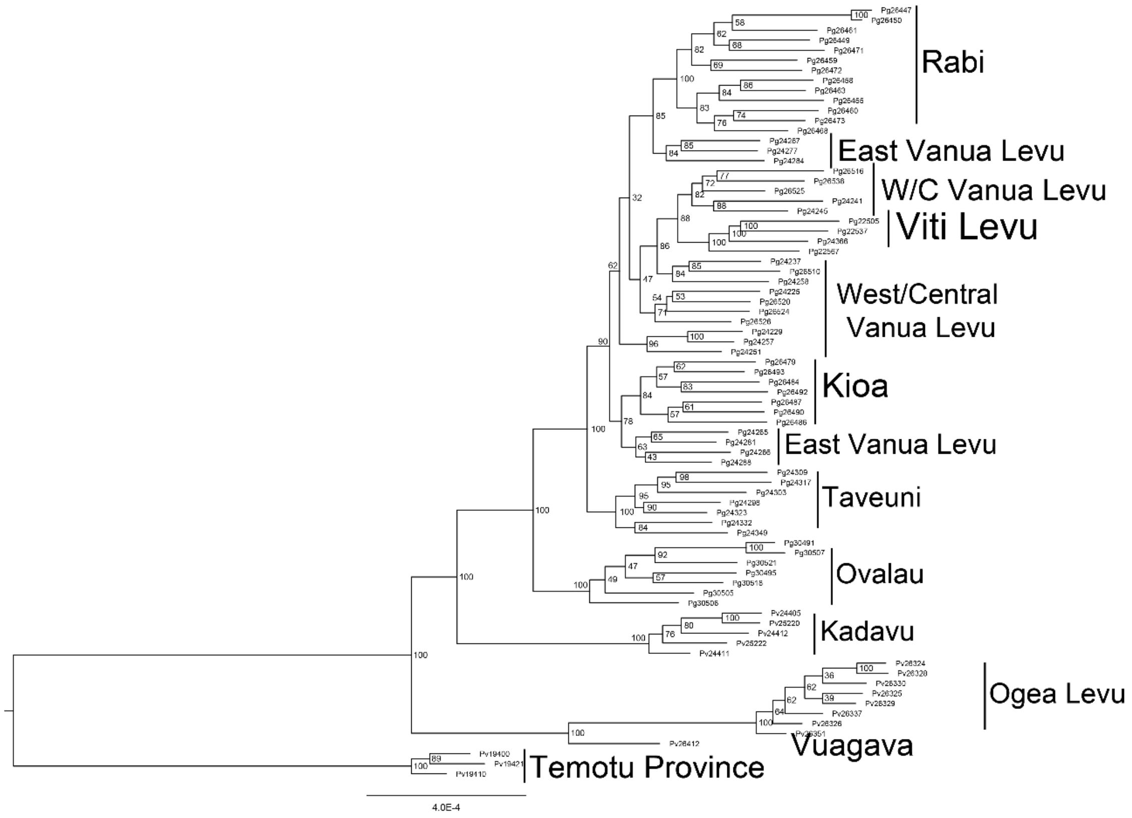

Figure S3: Full TreeMix analysis, as in Figure 2E, but including the network with only a single migration edge permitted. Tip names correspond to sampled populations, and inferred admixture proportion is labeled next to the arrows representing a given edge.

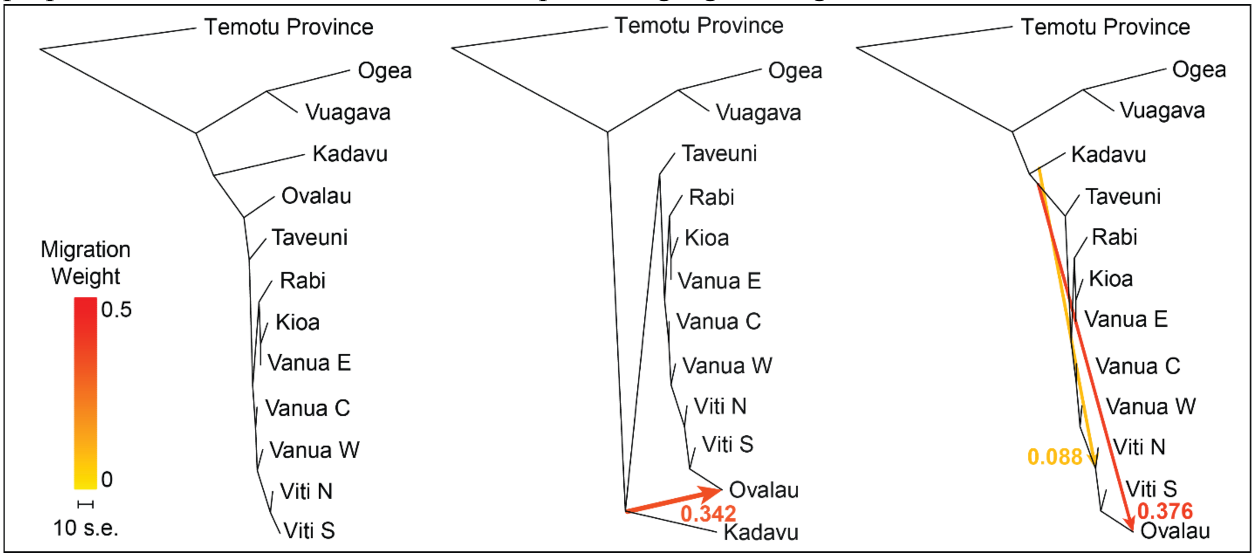


Figure S4: Comparison of two SVDQuartets cladograms, A) with and B) without Ovalau included. Bootstrap values are reported on nodes, and branches with values less than 70 are collapsed. Note that the inclusion of the admixed population on Ovalau confounds phylogenetic inference.

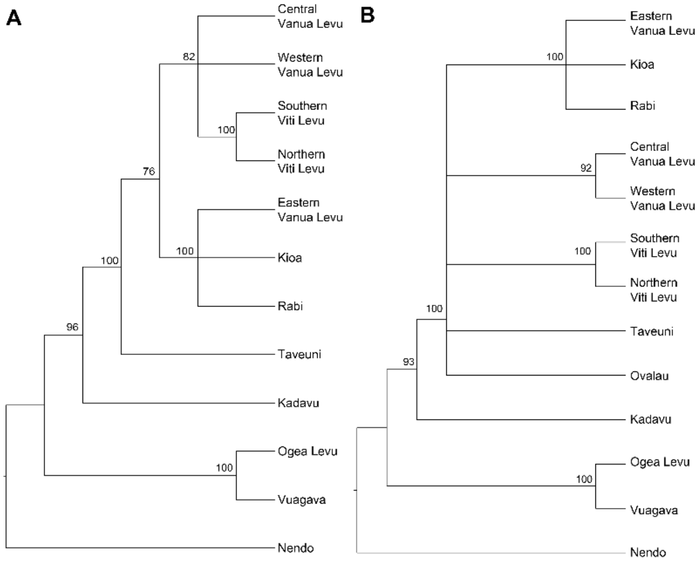


Figure S5: Time-calibrated phylogeny of ultraconserved element data for a wider sampling of *Pachycephala*, used in Figure 1. Posterior probabilities are displayed on nodes, and scale bar is in millions of years.

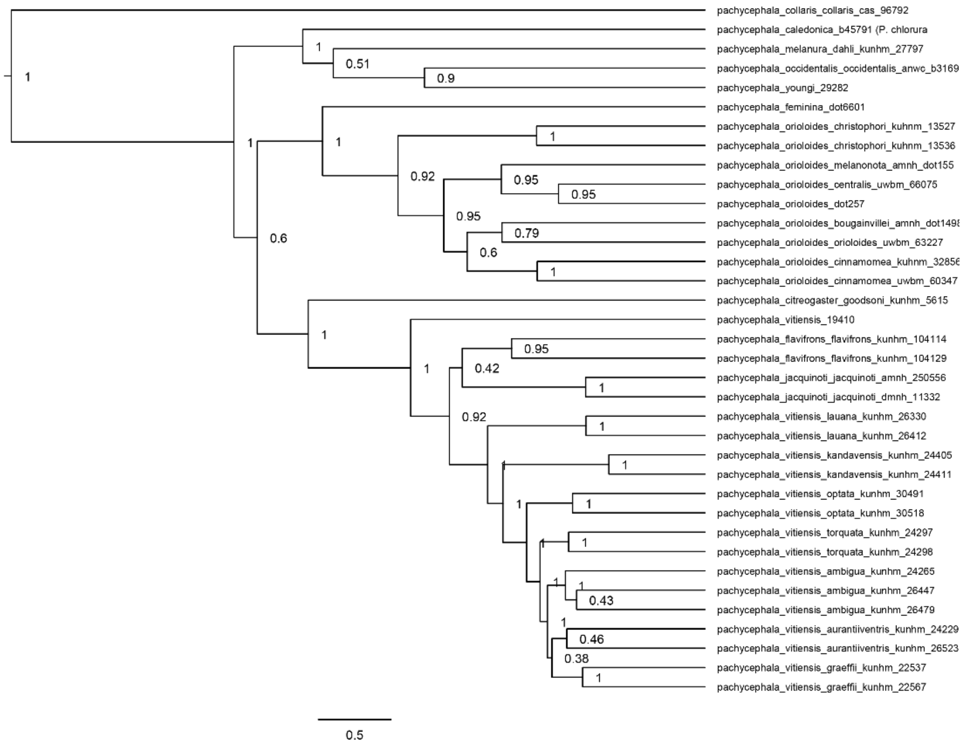


Figure S6: Plot representing significant values of the fbranch statistic, with the darkness of the colored squares representing the inferred value for a given node. Gray squares represent comparisons that could not be tested for gene flow. Axes have phylogenies that were tested, with dashed lines representing gene flow with ancestral nodes.

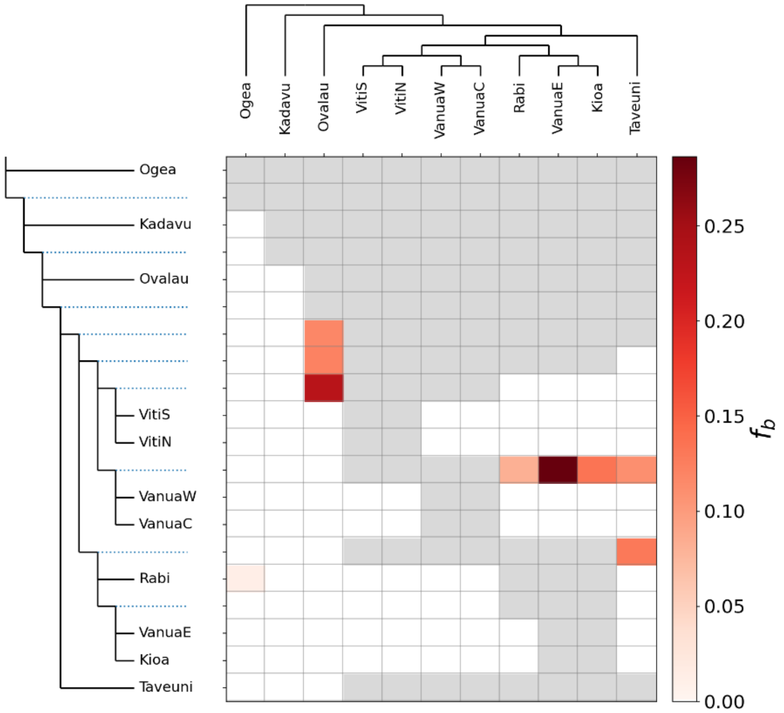

Figure S7: Additional Manhattan plots, as in Fig. 4. The Y-axis represents $F_{ST}$ in all comparisons, and the X-axis represents genome position. Titles describe the populations considered in a given comparison.

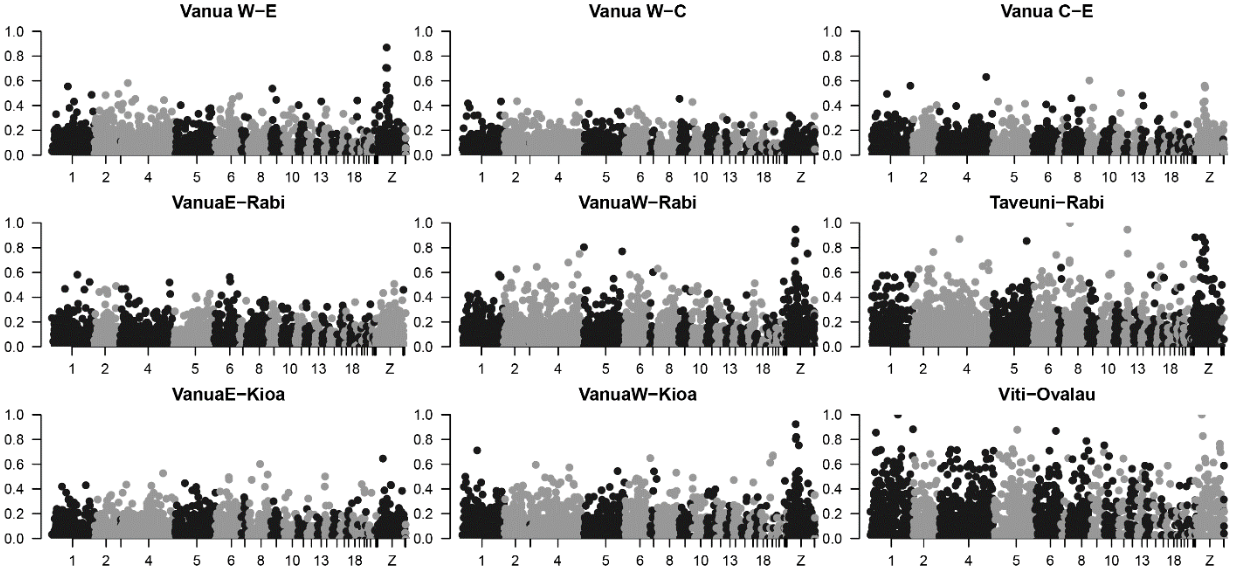


Table S1: Table of samples used for RAD-seq dataset. The columns from left to right are species, subspecies, sex, male phenotype scores, country, island (with locality in parentheses for larger islands), tissue number, the sample name used in scripts (note older taxonomy), and SRA sample name, number of reads, and dataset(s) in which the sample was used. Male phenotype was only scored for adult with study skins at the University of Kansas Biodiversity Institute (KU). Dataset numbers correspond to: 1) yellow-throated taxa only, 2) all taxa, 3) all taxa, phylogenetically confounding individuals removed individuals removed, and 4) no Ovalau, phylogenetically confounding individuals removed. **[IN DRYAD UPLOAD]**

Table S2: Table of samples used for UCE phylogeny and dating analysis. From left to right, columns are scientific name, country, island, institution, identifying number, source material (tissue vs historic toepad sample), sample name used in scripts, sample name used for SRA accession, NCBI accession number, sequencing platform, number of cleaned reads, number of recovered loci, and voucher number. **[IN DRYAD UPLOAD]**

Table S3: Table of autosome-wide (below diagonal) and Z-only (above diagonal) Weir and Cockerham $F_{ST}$ values. The diagonal is nucleotide diversity at all sites calculated by Stacks, with Viti Levu representing a combined value for north and south samples. Borders delimit major island groups in yellow-throated taxa. Column headers are abbreviated, but match their respective rows.

| Pops | Ova | VitiS | VitiN | VanW | VanC | VanE | Kioa | Rabi | Tav | Kad | Ogea | Nendo |
|---|---|---|---|---|---|---|---|---|---|---|---|---|
| Ovalau | 4.8E-04 | 0.249 | 0.251 | 0.202 | 0.171 | 0.221 | 0.248 | 0.255 | 0.280 | 0.400 | 0.653 | 0.553 |
| Viti S | 0.125 | 4.8E-04 | 0.033 | 0.133 | 0.129 | 0.234 | 0.269 | 0.298 | 0.325 | 0.518 | 0.777 | 0.712 |
| Viti N | 0.122 | 0.012 | 4.8E-04 | 0.077 | 0.085 | 0.228 | 0.288 | 0.302 | 0.338 | 0.518 | 0.799 | 0.736 |

| | | | | | | | | | | | | |
|---|---|---|---|---|---|---|---|---|---|---|---|---|
| Vanua W | 0.131 | 0.055 | 0.037 | 5.3E-04 | 0.016 | 0.087 | 0.117 | 0.147 | 0.212 | 0.366 | 0.631 | 0.525 |
| Vanua C | 0.120 | 0.057 | 0.035 | 0.016 | 5.5E-04 | 0.047 | 0.070 | 0.101 | 0.165 | 0.344 | 0.617 | 0.511 |
| Vanua E | 0.135 | 0.073 | 0.054 | 0.032 | 0.022 | 5.4E-04 | 0.005 | 0.049 | 0.147 | 0.367 | 0.630 | 0.532 |
| Kioa | 0.154 | 0.104 | 0.071 | 0.057 | 0.042 | 0.020 | 5.4E-04 | 0.063 | 0.156 | 0.393 | 0.661 | 0.574 |
| Rabi | 0.172 | 0.116 | 0.096 | 0.069 | 0.063 | 0.044 | 0.063 | 5.1E-04 | 0.190 | 0.391 | 0.631 | 0.551 |
| Taveuni | 0.171 | 0.121 | 0.102 | 0.079 | 0.067 | 0.064 | 0.078 | 0.094 | 5.3E-04 | 0.410 | 0.663 | 0.576 |
| Kadavu | 0.362 | 0.414 | 0.405 | 0.331 | 0.320 | 0.318 | 0.333 | 0.343 | 0.336 | 3.9E-04 | 0.753 | 0.686 |
| Ogea | 0.514 | 0.597 | 0.589 | 0.482 | 0.465 | 0.457 | 0.477 | 0.459 | 0.484 | 0.595 | 2.7E-04 | 0.843 |
| Nendo | 0.576 | 0.652 | 0.648 | 0.556 | 0.544 | 0.539 | 0.556 | 0.553 | 0.554 | 0.667 | 0.748 | 2.8E-04 |

Table S4: Summary of tests for multiple comparisons in ABBA/BABA tests, per each category. Columns are the category of edge, the number of tests, the harmonic p-value for the category, if it was significant before and after the Benjamini-Hochberg procedure, and newick notation of comparisons with significant tests in (P1, P2), P3 order.

| Category | Tests | Harmonic p-value | Significant before B-H? | Significant after B-H? | Comparisons with significant B-H |
|---|---|---|---|---|---|
| Kadavu - Other | 28 | 0.12904 | Yes | No | n/a |
| Lau - Other | 36 | 0.11439 | Yes | No | n/a |
| Taveuni - Other | 21 | 0.00616 | Yes | Yes | ((Viti S, Rabi), Taveuni); ((Viti N, Rabi), Taveuni); ((Viti N, Kioa), Taveuni); ((Viti S, Kioa), Taveuni) |
| Kioa - West | 6 | 0.01718 | Yes | Yes | ((Viti S, VanuaC), Kioa); ((Viti S, VanuaW), Kioa) |
| Rabi - West | 7 | 0.06841 | Yes | No | n/a |
| Vanua - East | 8 | 0.22257 | No | No | n/a |
| East - West Vanua | 6 | 0.05276 | Yes | No | n/a |
| Viti Levu - Other | 8 | 0.17925 | No | No | n/a |

Table S5: Detailed values for significant ($p<0.05$) ABBA/BABA tests before accounting for multiple comparisons, including the Ovalau population. It includes the D statistic, p-value associated with those D-statistics, f4 estimates of admixture, and category for multiple-comparison tests. Gene flow is between P2 and P3, but this approach does not test directionality of gene flow.

| P1 | P2 | P3 | D stat. | p-value | f4 | Category |
|---|---|---|---|---|---|---|
| VanuaE | VanuaC | Kadavu | 0.0582 | 0.0403 | 0.0305 | Kadavu - Other |
| VanuaE | VitiN | Kadavu | 0.0708 | 0.0463 | 0.0367 | Kadavu - Other |

| | | | | | | |
|---|---|---|---|---|---|---|
| VanuaE | VitiS | Kadavu | 0.0801 | 0.0300 | 0.0429 | Kadavu - Other |
| VitiS | VanuaC | Kioa | 0.0746 | 0.0041 | 0.3216 | Kioa - West |
| VitiS | VanuaW | Kioa | 0.0620 | 0.0157 | 0.2704 | Kioa - West |
| Ovalau | VitiS | Kioa | 0.1798 | 0.0000 | 0.4577 | n/a |
| Ovalau | Kioa | Ogea | 0.0962 | 0.0168 | 0.0322 | n/a |
| Ovalau | Rabi | Ogea | 0.1203 | 0.0034 | 0.0401 | n/a |
| VanuaE | Rabi | Ogea | 0.0816 | 0.0135 | 0.0248 | Lau - Other |
| Taveuni | Rabi | Ogea | 0.0697 | 0.0405 | 0.0213 | Lau - Other |
| Ovalau | VanuaC | Ogea | 0.1046 | 0.0084 | 0.0325 | n/a |
| Ovalau | VitiN | Ogea | 0.0905 | 0.0352 | 0.0275 | n/a |
| Ovalau | VitiS | Ogea | 0.1165 | 0.0123 | 0.0352 | n/a |
| Taveuni | Kioa | Ovalau | 0.0770 | 0.0040 | 0.1186 | n/a |
| Taveuni | Rabi | Ovalau | 0.0893 | 0.0007 | 0.1368 | n/a |
| Taveuni | VanuaC | Ovalau | 0.1560 | 0.0000 | 0.2500 | n/a |
| Kioa | VanuaC | Ovalau | 0.0847 | 0.0005 | 0.1477 | n/a |
| VanuaE | VanuaC | Ovalau | 0.0685 | 0.0024 | 0.1280 | n/a |
| Rabi | VanuaC | Ovalau | 0.0705 | 0.0031 | 0.1230 | n/a |
| Taveuni | VanuaE | Ovalau | 0.0907 | 0.0004 | 0.1382 | n/a |
| Taveuni | VanuaW | Ovalau | 0.1561 | 0.0000 | 0.2451 | n/a |
| Kioa | VanuaW | Ovalau | 0.0855 | 0.0009 | 0.1571 | n/a |
| Rabi | VanuaW | Ovalau | 0.0720 | 0.0032 | 0.1327 | n/a |
| VanuaE | VanuaW | Ovalau | 0.0698 | 0.0036 | 0.1225 | n/a |
| Kioa | VitiN | Ovalau | 0.1870 | 0.0000 | 0.3506 | n/a |
| VanuaE | VitiN | Ovalau | 0.1716 | 0.0000 | 0.3380 | n/a |
| Rabi | VitiN | Ovalau | 0.1734 | 0.0000 | 0.3172 | n/a |
| VanuaC | VitiN | Ovalau | 0.1087 | 0.0003 | 0.2366 | n/a |
| VanuaW | VitiN | Ovalau | 0.1106 | 0.0003 | 0.2285 | n/a |
| VanuaW | VitiS | Ovalau | 0.1567 | 0.0000 | 0.3481 | n/a |
| VanuaC | VitiS | Ovalau | 0.1544 | 0.0000 | 0.3562 | n/a |
| VitiN | VanuaC | Rabi | 0.0502 | 0.0357 | 0.1637 | Rabi - West |
| VitiN | VanuaW | Rabi | 0.0527 | 0.0333 | 0.1724 | Rabi - West |
| VitiS | VanuaW | Rabi | 0.0481 | 0.0462 | 0.1667 | Rabi - West |
| Ovalau | VitiS | Rabi | 0.2070 | 0.0000 | 0.4198 | n/a |
| VitiN | Kioa | Taveuni | 0.0782 | 0.0041 | 0.3575 | Taveuni - Other |
| VitiS | Kioa | Taveuni | 0.0758 | 0.0050 | 0.3306 | Taveuni - Other |
| VitiS | Rabi | Taveuni | 0.0935 | 0.0008 | 0.3798 | Taveuni - Other |
| VitiN | Rabi | Taveuni | 0.0940 | 0.0008 | 0.3891 | Taveuni - Other |
| VanuaW | Rabi | Taveuni | 0.0459 | 0.0328 | 0.2218 | Taveuni - Other |
| VitiN | VanuaC | Taveuni | 0.0602 | 0.0229 | 0.2591 | Taveuni - Other |
| VitiS | VanuaC | Taveuni | 0.059 | 0.0251 | 0.2454 | Taveuni - Other |
| VitiS | VanuaE | Taveuni | 0.063 | 0.0145 | 0.2622 | Taveuni - Other |
| VitiN | VanuaE | Taveuni | 0.062 | 0.0168 | 0.2609 | Taveuni - Other |
| VitiS | VanuaW | Taveuni | 0.052 | 0.0452 | 0.2237 | Taveuni - Other |

| | | | | | | |
|---|---|---|---|---|---|---|
| Ovalau | VitiN | Taveuni | 0.168 | 6E-07 | 0.4046 | n/a |
| Ovalau | VitiS | Taveuni | 0.164 | 1E-06 | 0.3872 | n/a |
| VitiN | VanuaC | VanuaE | 0.053 | 0.0243 | 0.582 | East - West Vanua |
| VitiS | VanuaC | VanuaE | 0.048 | 0.0388 | 0.5265 | East - West Vanua |
| VitiN | VanuaW | VanuaE | 0.047 | 0.0441 | 0.5727 | East - West Vanua |
| Ovalau | VitiS | VanuaE | 0.192 | 2E-09 | 0.6834 | n/a |
